# Striped distribution pattern of Purkinje cells of different birthdates in the mouse cerebellar cortex studied with the *Neurog2*-CreER transgenic line

**DOI:** 10.1101/2020.05.10.086603

**Authors:** Jingyun Zhang, Khoa Tran-Anh, Tatsumi Hirata, Izumi Sugihara

## Abstract

Heterogeneity of Purkinje cells (PCs) that are arranged into discrete longitudinal stripes in the cerebellar cortex is related to the timing of PC generation. To understand the cerebellar compartmental organization, we mapped the PC birthdate (or differentiation timing) in the entire cerebellar cortex. We used the birthdate-tagging system of *neurog2*-CreER (G2A) mice hybridized with the AldocV strain which clarifies the zebrin (aldolase C) longitudinal striped pattern.

The pattern of the birthdate-dependent PC distribution was arranged consistently into longitudinally-oriented stripes throughout almost all lobules except for the nodulus, paraflocculus and flocculus, in which distinct stripes were observed.Boundaries of the PC birthdate stripes were found either in the middle or coincided with that of the zebrin stripes. PCs in each birthdate stripe were born in various periods between embryonic day (E) 10.0 and E 13.5.

In the vermis, PCs were chronologically distributed from lateral to medial stripes. In the paravermis, PCs of early birthdates were distributed in the long lateral zebrin-positive stripe (stripe 4+//5+) and the medially neighboring narrow zebrin-negative substripe (3d-//e2-), while PCs of late birthdates were distributed in the rest of all paravermal areas. In the hemisphere, PCs of early and late birthdates were intermingled in the majority of areas.

The results indicate that the birthdate of a PC is a partial determinant for the zebrin compartment in which it is located. However, the correlation between the PC birthdate and the zebrin compartmentalization is not simple, and distinct among the vermis, paravermis, hemisphere, nodulus, and flocculus.

**Highlights:** Birthdates of Purkinje cells (PCs) were mapped on the cerebellar zebrin striped pattern by using *Neurog2*-CreER (G2A) mice.

The vermis, paravermis, hemisphere, nodulus, and flocculus had distinct longitudinally-striped patterns of PC birthdate distribution.

PCs in each birthdate stripe were born in various periods between embryonic day (E) 10.0 and E 13.5.

Boundaries of PC birthdate distributions were located at the boundaries of zebrin stripes or in the middle of a zebrin stripe.

The results indicate that the PC birthdate is a partial determinant for the zebrin compartment in which a PC is located.

## 1 Introduction

The cerebellar cortex is conventionally subdivided into three major subdivisions; vermis, paravermis, and hemisphere, which are connected with the medial, interposed, and lateral cerebellar nuclei via axonal projections of Purkinje cells (PCs) - the sole output neurons of the cerebellar cortex (Ito, 1984). Clarification of some details of efferent PC axons and afferent climbing fiber axons has led to the definition of more detailed subdivisions named basically by alphanumeric characters: A, lateral A, B, X, C1, C2, C3, CX, D0, D1 and D2 modules (Ruigrok et al., 2015; Buissert-Delmas et al, 1993; Voogd and Bigaré, 1980). Among these modules, module B, which project to the dorsal part of the lateral vestibular nucleus also called Deiters’ nucleus, was used to demonstrate the inhibitory nature of PCs by Ito and Yoshida (1966)

Later, another type of subdivisions has been defined by longitudinally-striped compartments of PCs that have common expression profile of several molecules including aldolase C (zebrin) (Brochu et al., 1990; Voogd et al., 2003; Sugihara and Shinoda, 2004; Fujita et al., 2014). Since the positional relationship between zebrin-based subdivisions and projection-based modules has been identified, the zebrin striped pattern can be used as the standard reference map of the rodent cerebellum. Expression patterns of various molecules, mossy fiber afferent projection patterns, and neuronal responsiveness have been mapped on the zebrin striped pattern to understand the morphological and functional organization of the cerebellum (Voogd et al., 2003; Sugihara and Shinoda, 2004; Sugihara et al., 2009; Apps and Hawkes, 2009; Quy et al., 2011; Tsutsumi et al., 2019; Luo et al., 2019).

PCs are born during embryonic days (E)10.5–12.5 in the ventricular zone (Hashimoto and Mikoshiba, 2003) and arranged into clusters separated by PC-free gaps in the three-dimensional (3D) space in the superficial part of the cerebellum by E17.5 (Korneliussen, 1968; Altman and Bayer, 1997; Wassef and Sotelo, 1984; Smeyne et al., 1991; Oberdick et al., 1993; Millen et al., 1995; Larouche et al., 2006; Wilson et al., 2011; Fujita et al., 2012) in the mouse. These clusters of PC subsets are the direct origin of longitudinally striped compartments of the adult cerebellar cortex (Sillitoe et al., 2009, 2010; Namba et al., 2011; Fujita et al., 2012; Vibulyaseck et al., 2017).

Previous studies with replication-defective adenoviral vector labeling of newly generated PCs showed that PCs born at a particular birthdate (E10.5, E11.5, or E12.5) are distributed in a specific set of zebrin stripes (Hashimoto and Mikoshiba, 2003). Early (E10.5-born; Namba et al., 2011) PCs were distributed in the lateral zebrin-positive (Z+) stripe in the vermis, paravermis, and hemisphere. Intermediate (E11.5-born) PCs were distributed in more medial Z+ and zebrin-negative (Z-) stripes in the vermis, paravermis, and hemisphere. Late (E12.5-born) PCs were distributed in the most medial Z+ and Z-stripes of the vermis, paravermis, and hemisphere (Namba et al., 2011). Thereby, it has been proposed that PC birthdate directly controls the fate of PCs and the similar birthdate-dependent organization shared by the vermis, paravermis, and hemisphere (Namba et al., 2011).

Recently a different efficient method for identifying the neuronal birthdate with the CreER-LoxP system involving a specific gene that is transiently expressed around the time of neuronal birth has been developed (Joyner and Zervas, 2006). Indeed, this method performed in the cerebellum with *Ascl1* gene, which is expressed at the transition from proliferating neuronal progenitors to differentiating neurons, has shown that PCs labeled by tamoxifen given at E10.5, E11.5, and E12.5 are distributed differently in the adult cerebellum (Sudarov et al., 2011). In the present study, we employed a similar method of neuronal birthdate tagging with *Neurog2*-CreER (G2A) mice (Hirata et al., 2019). The *Neurog2* gene is expressed transiently in some neurons including PCs after the final mitosis of neurogenesis (Florio et al., 2012). By giving tamoxifen at various 12-hour interval timing from E9.5 to E15.5 to the pregnant mother of G2A mice hybridized with loxP reporter (Ai9) and *Aldoc* reporter (AldocV) strains, we analyzed the distribution of PCs of different birthdates systematically on the striped zebrin expression pattern in the entire cerebellar cortex. Our results clarified the detailed positional relationship between the pattern of birthdate-dependent distribution of PCs and the zebrin (aldolase C) striped pattern in the adult mouse cerebellar cortex.

## 2 Materials and Methods

### 2.1 Ethics statements

Experimental protocols were approved by the Animal Care and Use Committee (A2019-187A, A2018-148A, A2017-060C4) and Gene Recombination Experiment Safety Committee (G2019-020A, 2017-040A, 2012-064C4) of Tokyo Medical and Dental University.

### 2.2 Animals

Mice were bred and reared in a 12-12 hour light-dark cycled condition in the animal facility of the university with freely available food and water. The C57BL/6N-Tg(Neurogenin2-CreER) mouse strain (G2A, CDB:0512T-1, Hirata et al., 2019) has the transgene putatively in the Y chromosome. G2A male mice were mated with C57BL/6N female to obtain the next generation of G2A mice. Genotype was checked in the tail sample by polymerase chain reaction with the primer for the Cre gene (Aki553, TAAAGATATCTCACGTACTGACGGTG, and Aki554, TCTCTGACCAGAGTCATCCTTAGC). In G2A mice, the tamoxifen-inducible CreER gene has replaced the coding sequence of the neurogenin2 gene (*neurog2*) in a genomic BAC transgene (Hirata et al., 2019). Since the CreER is expressed in differentiating PCs after the last mitosis under the *neurog2* enhancer (Florio et al., 2012), a tamoxifen administration at a certain developmental stage induces Cre recombination activity only in the PCs that are shortly after the last mitosis. As for the reporter mice, we produced double homo hybrid mice (Ai9::AldocV) from Aldoc-Venus:C57BL/6N strain (Fujita et al., 2014) and Ai9:57BL/6J strain (B6.Cg-Gt(ROSA)26Sor^tm9(CAG-tdTomato)Hze^/J, The Jackson Laboratory, https://www.jax.org/strain/007909). Aldoc-Venus mice express yellow-green fluorescent protein Venus in place of aldolase C (Aldoc), which is expressed in the striped pattern in the cerebellum, while Ai9 mice express red fluorescent protein tdTomato in the Cre-dependent manner. Ai9::AldocV double homo females were mated with G2A males. The day when the vaginal plug was detected in mating was designated as E0.5. Tamoxifen (T5648-1G, Sigma, St. Louis, MO, U.S.A.) was dissolved in corn oil (9 mmole/l, 032-17016, Wako, Wako Pure Chemical Industries, Ltd., Osaka, Japan). Tamoxifen was injected intraperitoneally (2.25 µmole/mouse) one time at noon when the embryos were E9.5, E10.5, E11.5, E12.5, E13.5 or E14.5, or at midnight when the embryos were E10.0, E11.0, E12.0 or E13.0. The pregnant females were sacrificed by cervical dislocation to take out embryos with the Cesarean section when embryos were E19.5. Mice were reared by a stepmother. Control mice were born from the female in which tamoxifen was not given and were reared by their mother. Female pups were discarded in the first postnatal week. At postnatal day 40-42, male pups (designated as G2A::Ai9::AldocV mice in this study) were overdosed with an intraperitoneal injection of pentobarbital sodium (0.18 mg/g body weight) and xylazine (0.009 mg/g body weight) and perfused transcardially with phosphate-buffered saline (PBS, pH=7.4) with heparin sulfate (0.1%), and then with 4% paraformaldehyde. The brain and the spinal cord were dissected after over-night postfixation in 4% paraformaldehyde and soaked in 30% sucrose with phosphate buffer (pH=7.4) for two days. Brain samples were stored in the deep freezer before cutting.

To change the dose of tamoxifen, 1/2, 1/4, or 1/8 of the above tamoxifen amount was injected. To obtain control samples, no tamoxifen was given (no tamoxifen control), or a wild type C57BL/7N male mouse was mated with an Ai9::AldocV double-homo female in place of the G2A male mouse (Ai9::AldocV mice).

### 2.3 Histological procedures

Brains were coated with gelatin solution (10% gelatin, 10% sucrose in 10 mM phosphate buffer, 32 °C). The gelatin block was hardened by chilling and then soaked overnight in fixative with a high sucrose content (4% paraformaldehyde, 30% sucrose in 0.05 M phosphate buffer, pH 7.4). Serial coronal and horizontal sections were cut on a freezing microtome at a thickness of 50 µm and mounted on a slide. They are embedded with water-soluble mounting medium (CC mount, C9368-30ML, Sigma) and coverslipped.

### 2.4. Acquisition of digital images

Fluorescence images were digitized using a cooled color CCD camera (AxioCam1Cm1, Zeiss, Oberkochen, Germany) attached to a fluorescent microscope (AxioImager.Z2, Zeiss) in 16-bit gray-scale with an appropriate filter set. Serial coronal and horizontal sections of all experimental samples were digitized with a 2.5x objective, appropriate filter sets, and tiling function of the software (Zen, Zeiss). Several areas of particular sections were digitized with a higher magnification objective or with a confocal microscope (TCS SP8, Leica, Wetzlar, Germany) with a 40x objective. Images were adjusted concerning contrast and brightness and assembled using a software (Zen; Photoshop 7, Adobe, San Jose, CA, U.S.A.). An appropriate combination of pseudo-color was applied to fluorescent images. Photographs were assembled using Photoshop and Illustrator software (Adobe). The software was used to adjust contrast and brightness, but no other digital enhancements were applied.

### 2.5. Mapping tdTomato-labeled PCs on the scheme of the unfolded cerebellar cortex with the zebrin striped pattern

The striped zebrin (aldolase C) expression pattern has been drawn on the unfolded representation of the mouse cerebellar cortex (Sugihara and Quy, 2007; Fujita et al., 2014). The latest version of this scheme shown in Sarpong et al. (2018) was used. The nomenclature of zebrin stripes is complicated. For example, different names are used for the linked or equivalent stripes in rostral and caudal parts of the cerebellar cortex. To indicate such a linked pair of zebrin stripes, a double asterisk (“//”) is used (Sugihara and Shinoda, 2004). Nomenclature and definition of zebrin stripes have been described in our previous reports (Sugihara and Shinoda, 2004; Sugihara and Quy, 2007; Fujita et al., 2014; Sarpong et al 2018). To identify the location or the zebrin stripe where tdTomato-labeled PCs were located, we referred to the atlas of zebrin stripes in AldocV mice (Supplementary Figures 2–5 of Fujita et al., 2014). The zebrin stripe where tdTomato-labeled PCs were located was first identified by observing the green-fluorescence channel component of the original image in serial sections. Then, the identified stripes or substripes in which tdTomato-labeled PCs were observed were circumscribed on the scheme of zebrin stripes and indicated by the dotted pattern. The density of the tdTomato-labeled Purkinje cells was reflected in the density of the dots. Since we made multiple samples for most of the different tamoxifen injection timing, the averaged results were mapped in the scheme, although the interindividual difference was usually ignorable.

### 2.6. Serial section alignment analysis in the paraflocculus and flocculus

The method of serial section alignment analysis (SSAA) has been described (Fujita et al., 2012; Vibulyaseck et al., 2015; Sarpong et al., 2018). The PC and molecular layers in the rostral or caudal side of the paraflocculus and flocculus were clipped in digital images of serial horizontal sections. The clipped images were aligned in a position to clarify the striped pattern in a given area. The distribution pattern of tdTomato-labeled PCs was plotted on the unfolded scheme of paraflocculus and flocculus (Sarpong et al., 2018).

### 2.7. Analysis of PC birthdate range in Z+ stripes

The number of Z+ PCs in a given Z+ stripe was measured in the digital image of the section. Then the number of tdTomato-labeled PCs was measured among these Z+ PCs. This measurement was facilitated by turning on and off either the red+blue channel that represents the fluorescent signal of tdTomato or the green channel that represents the fluorescent signal of Venus in the digital image. Then the number of tdTomato-labeled PCs was divided by the number of all PCs to obtain “labeling percentage”. The same measurement was made for a given zebrin stripe in a given lobule in 4–12 sections at nearly the same rostrocaudal level in the cerebellums of various tamoxifen injection timing. The counted number of PCs for a Z+ stripe with each tamoxifen injection timing was between 33 and 333. Measured labeling percentage was plotted in the ordinate against the tamoxifen injection timing in the abscissa to in a graph to show the PC birthdate range.

## 3. Results

### 3.1. Labeling in G2A::Ai9::AldocV mice

G2A::Ai9::AldocV mice with or without a tamoxifen injection showed no apparent phenotypes in the behavior or size of the body or brain. A tamoxifen injection induced tdTomato-labeling of PCs (Fig. 1A). To find the best dose of tamoxifen to label PCs, the dose of tamoxifen injected at E11.5 was changed systematically from 2.25 µmole/mouse (Hirata et al., 2019), to 1/2, 1/4 and 1/8 of this dose. PCs were densely labeled in stripe 5+ and medially neighboring thin Z-stripe (e2-) in the paramedian lobule and copula pyramidis with tamoxifen 2.25 µmole/mouse (Fig. 1A). They were also labeled slightly less densely with 1/2 dose (Fig. 1B), further less densely with 1/4 dose (Fig. 1C), and only sparsely with 1/8 dose (Fig. 1D). The results indicated that PC labeling was dependent on tamoxifen and the effect of tamoxifen in labeling PCs was roughly saturating at 2.25 µmole/mouse. Therefore, we used this dose in all experiments.

**Figure 1.**
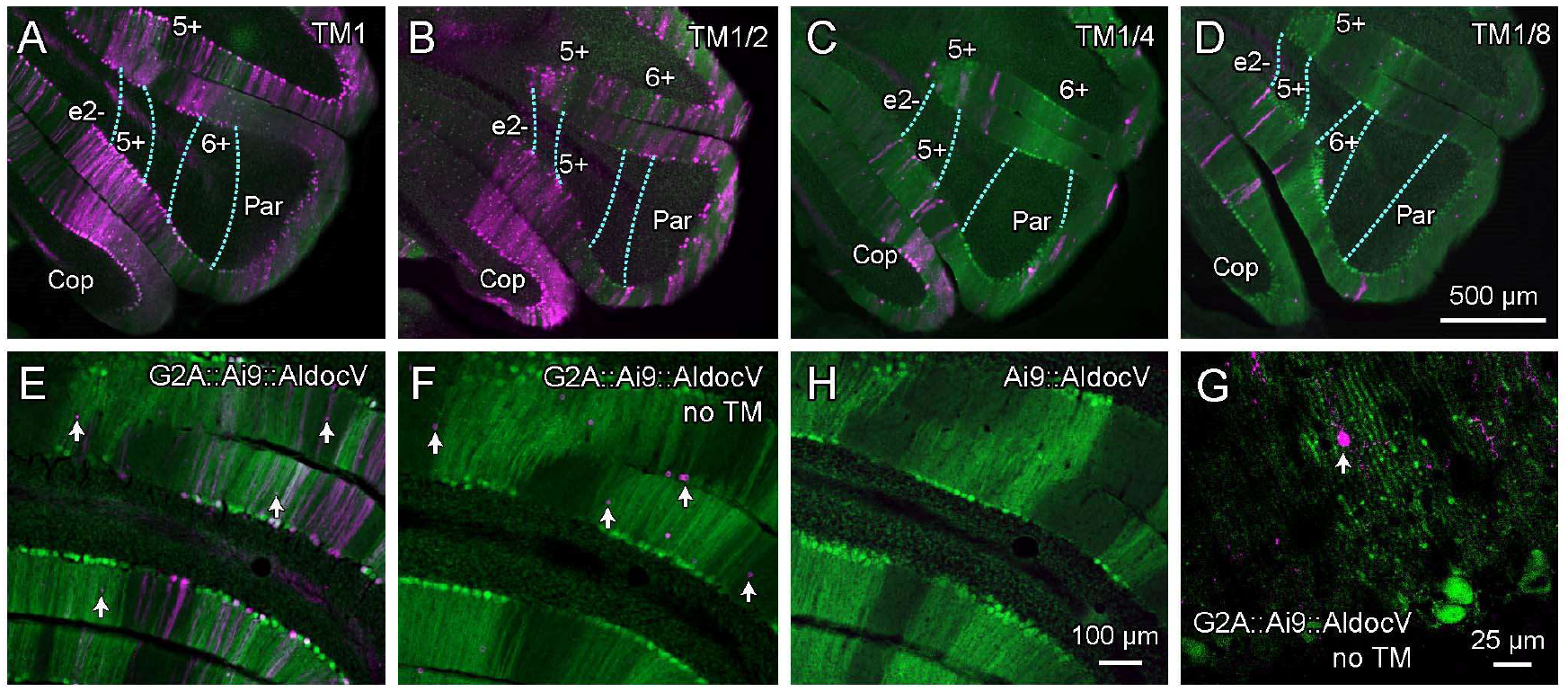
Efficiency and specificity of tamoxifen-dependent neuronal labeling in G2A::Ai9::AldocV mice. A-D, Images of the paravermal and hemispheric area of right crus II, paramedian lobule, and copula pyramidis in a coronal section of the G2A::Ai9::AldocV mice at P40–42. They were given a standard dose (2.25 µmole/mouse) (A), and 1/2 dose (B), 1/4 dose (C), and 1/8 dose (D) of tamoxifen at E11.5. E–G, More magnified image of the paravermal area of the paramedian lobule in the G2A::Ai9::AldocV mouse with an injection of the standard dose of tamoxifen at E11.5 (E), in the G2A::Ai9::AldocV mouse that was not given tamoxifen (F), and in the Ai9::AldocV mouse (G). Arrows in E indicate tdTomato-labeled cells that are not PCs. H, confocal images of tdTomato-labeled cells in the molecular layer in the G2A::Ai9::AldocV mouse that was not given tamoxifen. Scale bars, 500 µm in D (applies to A–D), 100 µm in G (applies to E–G), 25 µm in F.

**Figure 2.**
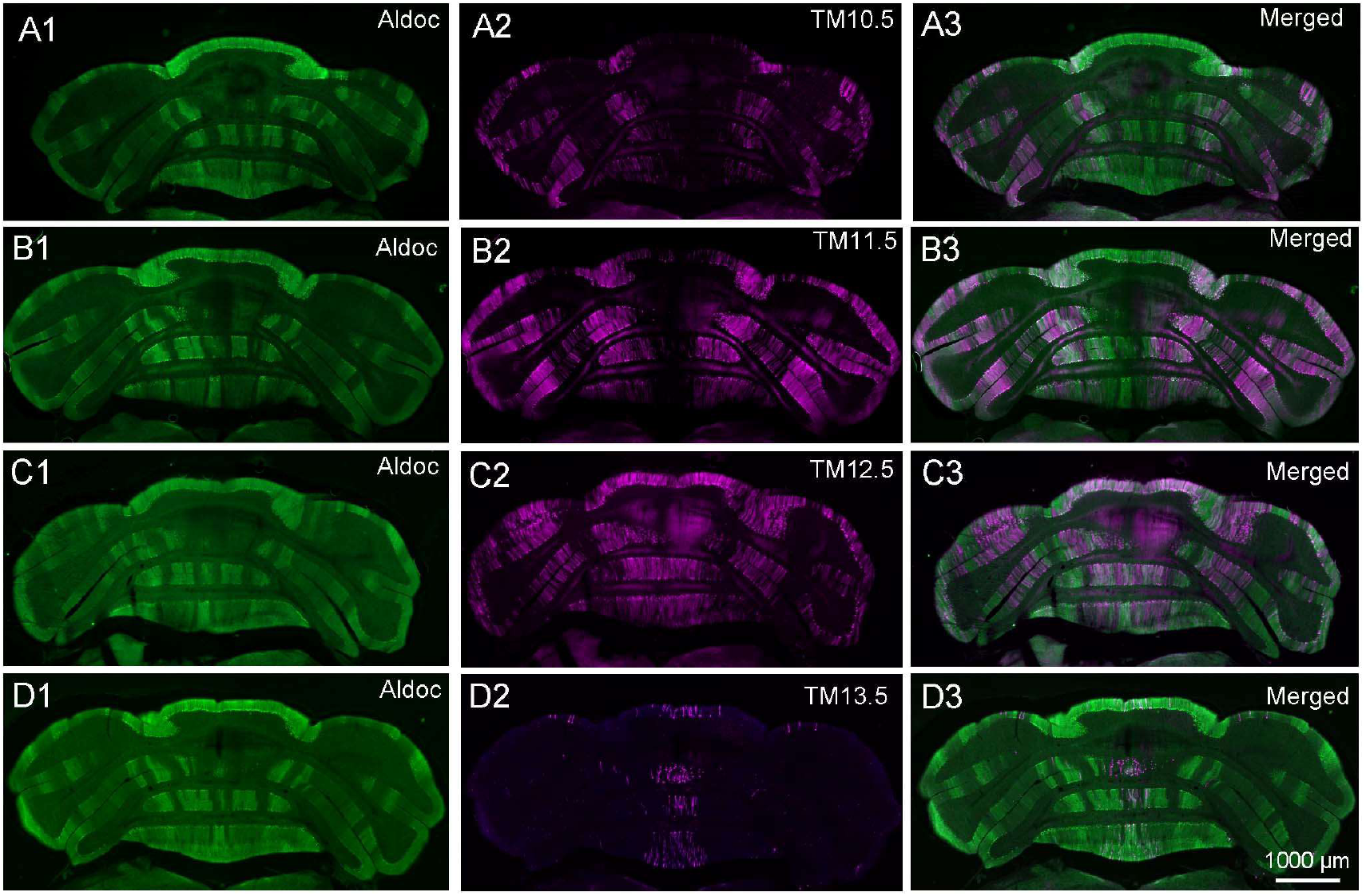
Overall distribution patterns of tdTomato-labeled PCs dependent on the timing of tamoxifen injection. A–D, Coronal sections of the caudal level of the cerebellum of G2A::Ai9::AldocV mice at P40–42. They were given tamoxifen at E10.5 (A), E11.5 (B), E12.5 (C) and E13.5 (D). Scale bars, 500 µm in A–D.

At higher magnification, we observed that small cells were also sparsely labeled in the molecular layer besides PCs (Fig. 1E, arrows). To examine the specificity of tamoxifen-dependent labeling of these cells, we compared labeling in control mice. In G2A::Ai9::AldocV mice without tamoxifen, small cells were still labeled with tdTomato sparsely in the molecular layer (Fig. 1F, arrows). Observation with confocal microscopy indicated they were a population of stellate cells in the molecular layer (Fig. 1H, arrow). In Ai9::AldocV mice, no cells were labeled with tdTomato (Fig.1G). Mechanisms of stellate cell labeling in G2A::Ai9::AldocV mice were not clear. Besides a population of stellate cells, labeling of PC-like cells occurred very rarely (not shown), only a few in the entire cerebellum in G2A::Ai9::AldocV mice without tamoxifen injection, which was ignorable. Therefore, we assumed that all PC labeling was induced by tamoxifen injection in the present study.

### 3.2. Overall fluorescent expression in the cerebellum in G2A::Ai9::AldocV mice

Tamoxifen injection (2.25 µmole/mouse) was made at E9.5 (n=1 dam), E10.0(n=1), E10.5 (n=2), E11.0 (n=1), E11.5 (n=2), E12.0 (n=2), E12.5 (n=3), E13.0 (n=1), E13.5 (n=2), E14.5 (n=1) or E15.5 (n=1) to label newly generated neurons at particular timing in G2A::Ai9::AldocV mouse embryos. The cerebellum in one or two samples obtained from each dam was cut in serial coronal sections. The cerebellum of another sample or two other samples obtained from each dam was cut in horizontal sections. In G2A::Ai9::AldocV mice, zebrin stripes were labeled with yellow-green fluorescence of Venus (shown in the green channel in figures) while birthdate-tagged PCs were labeled with red fluorescence of tdTomato (shown in the magenta or red+blue channel in figures). This double labeling allowed a detailed analysis of the distribution of birthdate-tagged PCs (Fig. 2).

PCs labeled with tamoxifen injection at different dates were distributed differently in the cerebellar cortex. With tamoxifen injection at E10.5 (designated as TM10.5), PCs were labeled in the lateral vermis, central paravermis, and the hemisphere (Fig. 2A). With TM11.5, areas of PC labeling became wider than that with TM10.5 (Fig. 2B). With TM12.5, areas that were negative with TM10.5 became positive, while the majority of the areas positive with TM10.5 became negative (Fig. 2C). Thus, the pattern became partially complementary to that with TM10.5. The midline areas in the vermis and a few other small areas stayed positive (Fig. 2D) with TM13.5. Thus, PCs labeled with tamoxifen injections at different dates distributed in distinct but partly overlapping patterns. These distribution patterns resemble those obtained by birthdate-dependent PC labeling with the replication-defective adenoviral vector (Namba et al., 2011) to some extent, indicating that the birthdate-tagging with G2A mice labeled PCs basically in a birthdate-dependent manner. Furthermore, the clear labeling pattern and high recombination efficiency of the birthdate tagging system with G2A mice seemed to facilitate a detailed analysis of the birthdate-dependent distribution pattern of PC in the entire cerebellar cortex in this study.

### 3.3. Landmark boundary with complementary distributions of birthdate-specific PCs

Analysis of the local relationship between the distribution pattern of tdTomato-labeled PCs and the zebrin striped pattern (Fig. 3) was essential in the present study. At the junction between vermal lobule VI and simple lobule, two clear Z+ stripes, 2b+, and 2b+s (2b+ satellite) were first identified in the image of the Venus fluorescence in coronal sections of TM10.5, TM11.5 and TM12.5 cerebellums (Fig. 3A1, B1, C1, respectively), by referring to the previous identification of zebrin stripes in the mouse cerebellar cortex (Sugihara and Quy, 2007; Fujita et al., 2014; Sarpong et al., 2018). In the image of tdTomato fluorescence, the dense distribution of labeled PCs with TM10.5 (designated as TM10.5 PCs) had a clear lateral boundary (arrowheads in Fig. 3A2). The merged image showed that the lateral clear boundary of TM10.5 PC distribution was located at the lateral edge of zebrin stripe 2b+ (arrowheads in Fig. 3A3). The same boundary was still recognized with TM11.5 although the presence of some tdTomato-labeled PCs in the area lateral to this boundary flawed its clarity (arrowheads in Fig. 3B2,3). Then, with TM12.5, the same boundary separated the distribution of tdTomato-labeled PCs again, with the dense distribution in the area lateral to it and absent distribution in the area medial to it (arrowheads in Fig. 3C2,3). In summary, distributions patterns of early (TM10.5 and TM11.5) and late (TM12.5) PCs showed not only a clear boundary at the same position (at the lateral boundary of zebrin stripe 2b+) but also a complimentary flip about this boundary. This type of clear boundaries (designated as “landmark boundaries”) were observed only in particular places in the cerebellum and were useful in describing the birthdate-dependent distribution pattern of PCs in the cerebellar cortex.

**Figure 3.**
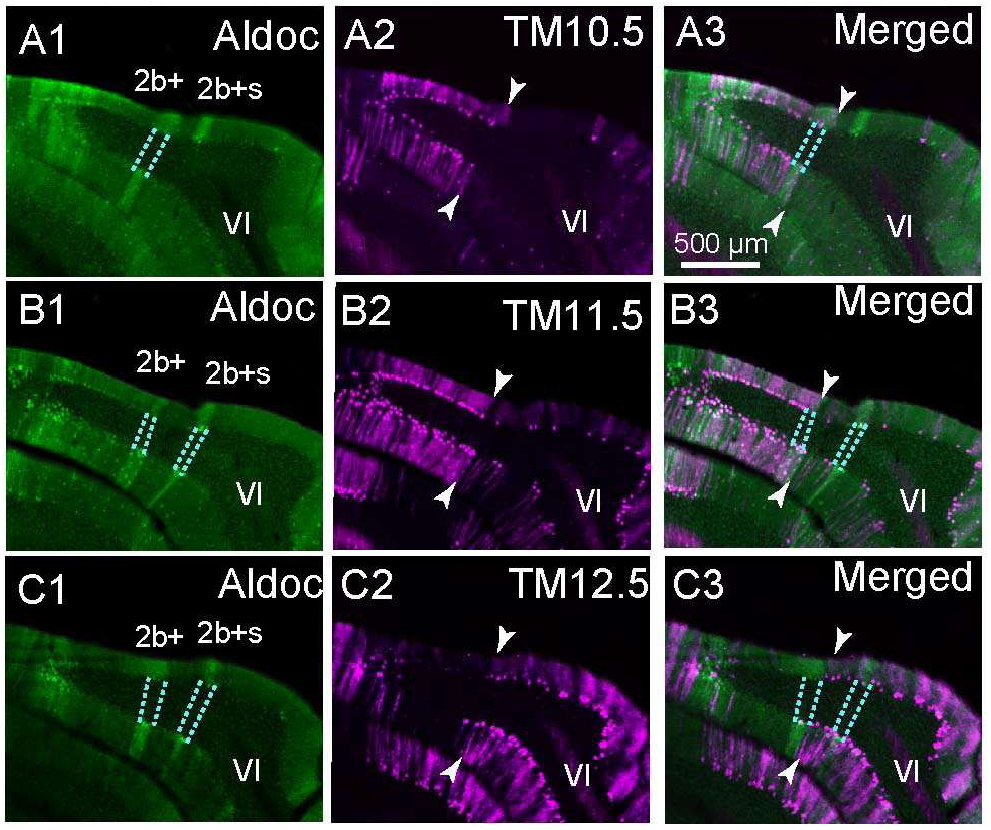
Local distribution patterns of tdTomato-labeled PCs showing a landmark boundary. E–G, Magnified coronal sections of lobule VI in the right cerebellum of G2A::Ai9::AldocV mice at P40-42. They were given tamoxifen at E10.5 (A), E11.5 (B), E12.5 (C). In each panel, the left, central, and right columns show the fluorescent signal of Venus, tdTomato, and both, respectively. Dashed lines indicate the boundaries of zebrin stripes. Arrowheads indicate a landmark boundary of tdTomato-labeled PCs. Scale bars, 100 µm in E–G.

### 3.4. Birthdate-dependent distribution pattern of PCs in vermal lobule IV-V

We then studied distribution patterns of labeled PCs in zebrin stripes systematically in several areas. In the vermal area of lobule IV-V, no PCs were labeled with TM9.5, TM14.5, or TM15.5 (Fig. 4A, K, L). Labeled PCs appeared in the medial portion of 2- and 2+, and very sparsely in the lateral part of 1-, with TM10.0 and TM10.5 (Fig. 4B, C). Labeled PCs still distributed in the medial portion of 2- and 2+, and additionally in the lateral portion of 1-with TM11.0, TM11.5, and TM12.0 (Fig. 4D–F).

**Figure 4.**
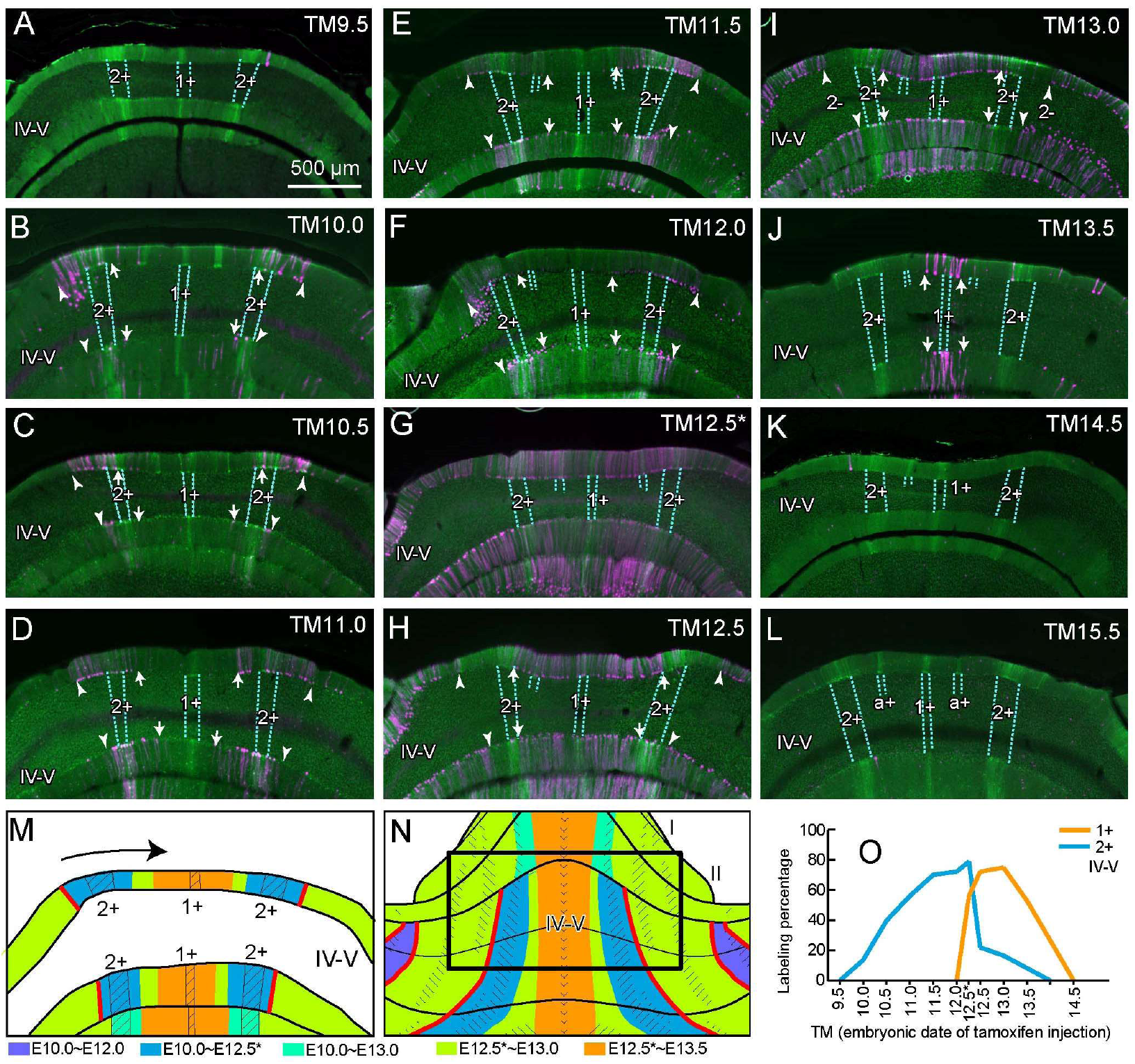
Distribution patterns of tdTomato-labeled PCs in vermal lobules IV-V in G2A::Ai9::AldocV mice at P40-42 with various tamoxifen injection timing. A-L, Images of the coronal cerebellar section at nearly the same level of G2A::Ai9::AldocV mice that were given tamoxifen at the timing indicated in the panel (E9.5–E15.5). Green and magenta channel shows the fluorescent signal of Venus and tdTomato, respectively. Arrowheads indicate the landmark boundary of the distribution of labeled PCs always located in the same position in stripe 2-. Arrows indicate other medial boundaries located in variable positions. Blue dashed lines indicate the boundaries between the Z+ and Z-stripes. TM12.5* indicates a particular case of tamoxifen injection at E12.5 which resulted in a labeling pattern of a slightly earlier stage. Scale bars, 500 µm in A-L, M. Schematic summary of PC birthdate distribution. Arrow indicates the chronological appearance of labeled PCs from the lateral to medial areas. N. Schematic summary of PC birthdate distribution in vermal lobules IV-V. Square indicates the approximate region shown in A–M. Shaded regions represent Z+ stripes in M and N. O, Analysis of PC birthdate range in stripes 1+ and 2+ of lobule IV-V with TM9.5– TM15.5 in panels A–L.

We obtained several mouse samples with TM12.5 from three dams. In mice obtained from one of the dams, PCs were labeled in almost all areas, less densely in 1+, medial 1-, 2+, and medial 2- (Fig. 4G). Since this pattern was partially different from the pattern observed in other mice with TM12.5, the mouse samples from this dam were designated as TM12.5*, while samples obtained from the two other dams were designated as TM12.5. In the TM12.5 and TM13.0 mice, PCs were labeled in all stripes except in 2+ and medial 2- (Fig. 4H, I). On the contrary, PCs were labeled in all stripes including 2+ and medial 2- in the TM12.5* mouse (Fig. 4G). This can be interpreted as the preservation of the TM11.5 pattern plus the appearance of the TM12.5 pattern. A possible reason for this was that the timing of tamoxifen injection was a little earlier than E12.5 for the TM12.5* embryos due to their incidental delayed development. PCs were labeled in 1+ and medial 1- with TM13.5 (Fig.4J). No PCs were labeled with TM14.5 or TM15.5 (Fig. 4K, L).

We also measured the density of labeled PCs with different tamoxifen injection timing in particular areas (PC birthdate range analysis). This analysis could be performed only in Z+ stripes, in which the total number of PCs could be counted thanks to PC labeling with Venus. Birthdates of PCs in stripe 2+ ranged broadly from TM10.0 to TM12.5, with a broad peak from TM11.0 to TM12.5* (Fig. 4O). In comparison, the birthdate distribution of PCs in stripe 1+ ranged narrowly from TM12.5* to TM13.5.

As a whole, the labeling pattern of PCs can be rather simply summarized. (1) Zebrin stripe 2- was subdivided into medial and lateral substripes that had different distributions of birthdate-specific PCs in lobule IV-V. (2) The boundary between the medial and lateral parts of zebrin stripe 2- was recognized as a landmark boundary (lateral arrowheads in Fig. 4B–F, H–I). A complete switching of the complementary labeling patterns occurs at this boundary within a short period between TM12.0 and TM12.5. (3) In the vermal area medial to this landmark boundary, PC labeling shifted from the lateral to medial areas along with the change of the tamoxifen injection timing from TM10.0 to TM13.5 (Fig. 4M, N; changing positions of arrows in Fig. 4B–J). Stripes 2+ and medial 2- were labeled with TM10.0–TM12.0; stripe lateral 1- were labeled with TM11.0–TM12.5; stripe a+ was labeled with TM12.5–TM13.0; stripe 1+ and medial 1- were labeled with TM12.5–TM13.5. Thus, no clear boundaries of birthdate-specific PC distribution were observed within the in the vermal areas medial to stripe 2-.

### 3.5. Birthdate-dependent distribution pattern of PCs in vermal lobule VIII

In the vermal area of lobule VIII, no PCs were labeled with TM9.5 (Fig. 5A). PCs were labeled sparsely in stripe 2- and moderately in stripes 3+, 3-, 4+ with TM10.0 and TM10.5 (Fig. 5B, C). Labeling persisted in the same stripes (2-, 3+, 3-, 4+) and started to appear in stripe 2-, the lateral part of stripe 2+ and stripes lateral to 4+ (i.e. stripes 4-, f+, f-, e1+, Sugihara and Shinoda, 2004; Fujita et al., 2014) with TM11.0, TM11.5, and TM12.0 (Fig. 5D-F). With TM12.5*, PC labeling nearly disappeared in stripes 3+ and 3- but it was observed in all other stripes (Fig. 5G). With TM12.5, PC labeling disappeared in stripes 3+, 3-, 4+ but persisted in all other stripes (Fig. 5H). With TM13.0, PC labeling also disappeared in lateral stripe 2+ and stripe 2- (Fig. 5I). With TM13.5, PC labeling stayed only in stripes 1+ and 1- (Fig. 5J). With TM14.5 and TM15.5, no PC labeling was observed (Fig. 5K, L).

**Figure 5.**
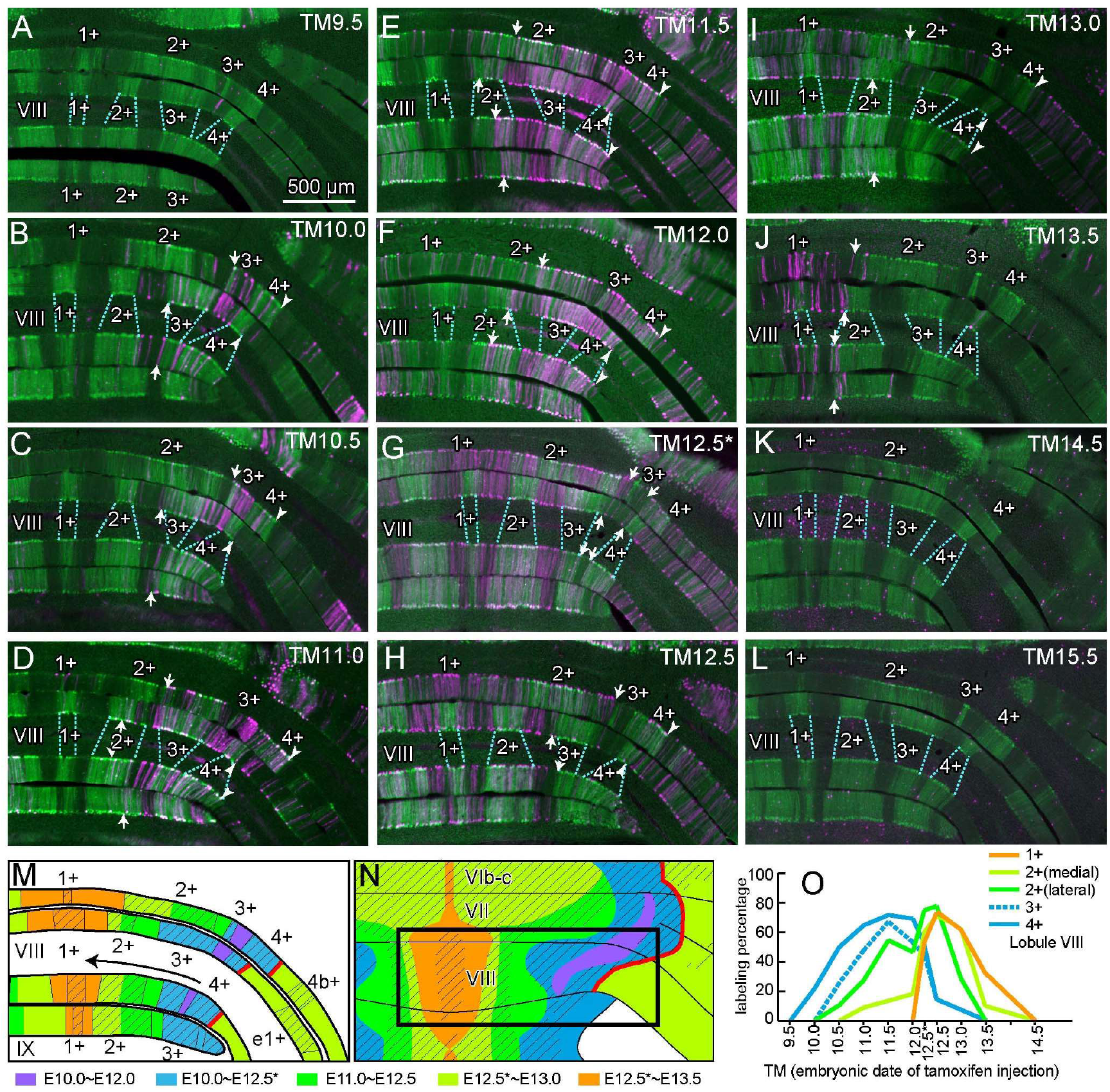
Distribution patterns of tdTomato-labeled PCs in vermal lobule VIII in G2A::Ai9::AldocV mice at P40-42 with various tamoxifen injection timing. A-L, Images of the coronal cerebellar section at nearly the same level of G2A::Ai9::AldocV mice that were given tamoxifen at the timing indicated in the panel (E9.5–E15.5). Green and magenta channel shows the fluorescent signal of Venus and tdTomato, respectively. Arrowheads indicate the landmark boundary of the distribution of labeled PCs at the lateral edge of stripe 4+. Arrows indicate other medial boundaries located in variable positions. Blue dashed lines indicate the boundaries between the Z+ and Z-stripes. Scale bars, 500 µm in A-L, M. Schematic summary of PC birthdate distribution. Arrow indicates the chronological appearance of labeled PCs from the lateral to medial areas. N. Schematic summary of PC birthdate distribution in vermal lobule VIII. Square indicates the approximate region shown in A–M. Shaded regions represent Z+ stripes in M and N. O, Analysis of PC birthdate range in stripes in stripes 1+, 2+, 3+, and 4+ of lobule VIII with TM9.5–TM15.5 in panels A–L.

In the PC birthdate range analysis, a gradual shift of the PC birthdate range was observed among different stripes (Fig. 5O). The timing-dependent increase of the labeling percentage was slower in early stripes but faster in late stripes (blue and green vs. yellow-green and orange in Fig. 5O).

As a whole, the labeling pattern of PCs can be rather simply summarized as follows. (1) Zebrin stripe 2+ in lobule VIII was subdivided into medial and lateral substripes that had different distributions of birthdate-specific PCs. (2) The boundary between stripes 4+ and 4- was recognized as the landmark boundary (lateral arrowheads in Fig. 5B–F, H–I). A complete switching of the complementary labeling pattern occurs at this boundary within a short period between TM12.0 and TM12.5. (3) In the vermal area medial to this landmark boundary, the PC labeling shifted from the lateral to medial areas along with the shift of the labeling timing from TM10.0 to TM13.5 (Fig. 5M, N; changing positions of arrows in Fig. 5B–J). Stripes 3+, 3- and 4+ were labeled with TM10.0–TM12.0; stripe lateral 2+ and 2- were labeled with TM11.0–TM12.5*; stripe medial 1+, 1- and medial 2+ were labeled with TM12.5*–TM13.0; stripe 1+ and medial 1- were still labeled with TM13.5. Thus, no clear boundaries of birthdate-specific PC distribution were observed within the vermal areas medial to stripe 2-. As we concluded in vermal lobules IV-V, our TM12.5* sample appeared to represent a little earlier in stage than other TM12.5 samples because it retained some aspects of the labeling pattern of TM12.0 (stripes 3+ and 4+).

### 3.6. Birthdate-dependent distribution pattern of PCs in paravermal lobule IV-V and simple lobule

In the paravermal lobule IV-V and simple lobule, no PCs were labeled with TM9.5 (Fig. 6A; a few PCs labeled in the most lateral part of 3- of lobule III). Many PCs were labeled in stripe 4+ and the most lateral part of 3- with TM10.0–TM12.0 (Fig. 6B–F). This most lateral Z- part between Z+ stripes 3+ and 4+ has been designated as 3b- because of the different olivocerebellar projection (Sugihara and Shinoda, 2004). This nomenclature was useful in the present study. Labeled PCs also appeared in the lateral part of 4-, 5+, and 5- in the same period (Fig. 6B–F). Then, labeled PCs were distributed in 3- except for the most lateral part (3b-), and in all stripes medial to 3-, 4-, and all stripes lateral to 4- with TM12.5* (Fig. 6G). Labeled PCs were distributed in 3- except for the most lateral part (3b-), and in all stripes medial to 3-, the medial part of 4- and 5- with TM12.5 and TM13.0 (Fig. 6H, I). No PCs were labeled with TM13.5, TM14.5, and TM15.5 (Fig. 6J, K, L).

**Figure 6.**
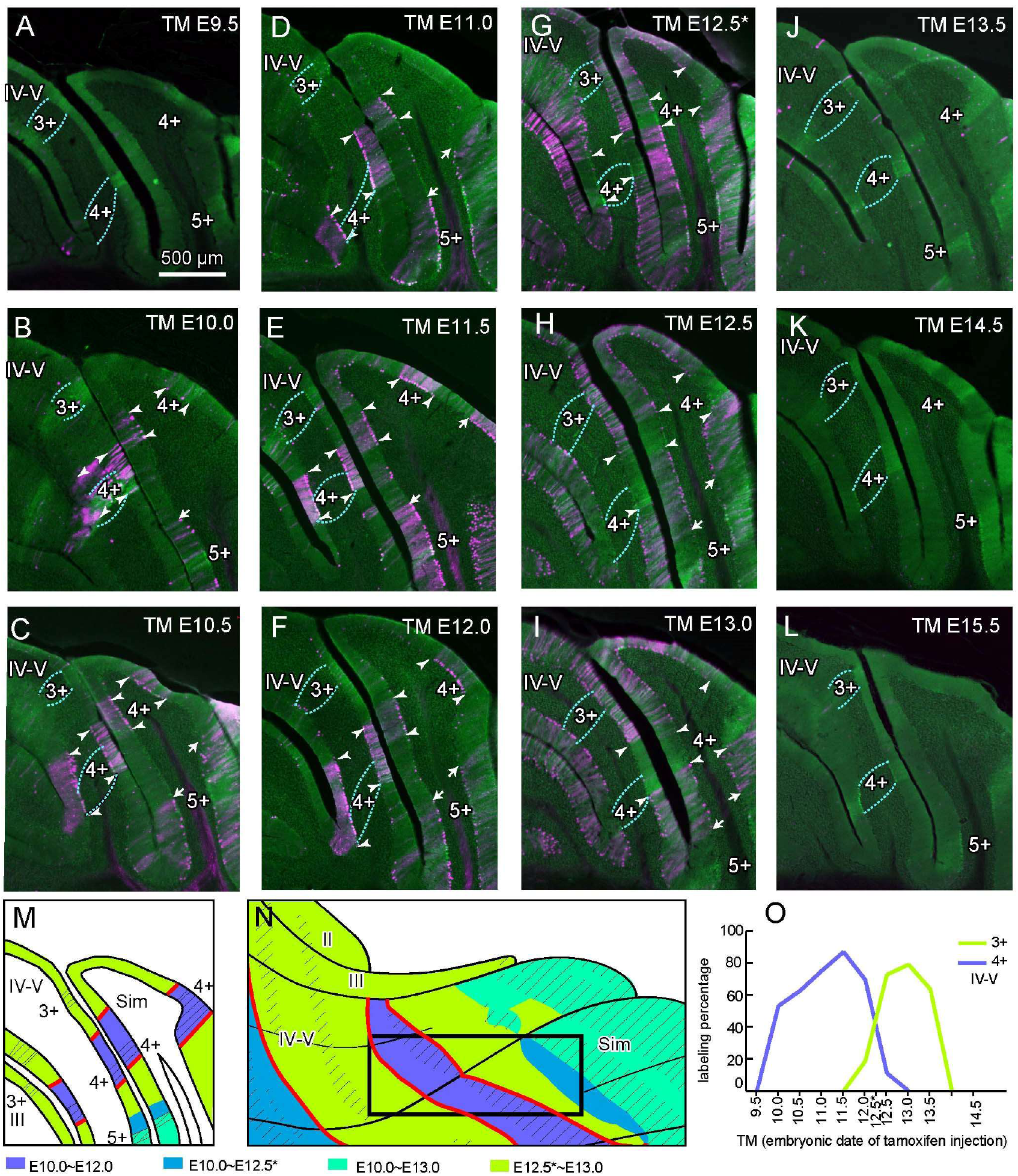
Distribution patterns of tdTomato-labeled PCs in paravermal lobules IV-V and simple lobule in G2A::Ai9::AldocV mice at P40–42 with various tamoxifen injection timing. A–L, Images of the coronal cerebellar section at nearly the same level of G2A::Ai9::AldocV mice that were given tamoxifen at the timing indicated in the panel (E9.5–E15.5). Green and magenta channels show the fluorescent signal of Venus and tdTomato, respectively. Arrowheads indicate the landmark boundaries of distribution of labeled PCs at the lateral edge of stripe 4+ and in the middle of stripe 3-. Arrows indicate other medial boundaries located in variable positions. Blue dashed lines indicate the boundaries between the Z+ and Z- stripes. Scale bars, 500 µm in A–L, M. Schematic summary of PC birthdate distribution. N. Schematic summary of PC birthdate distribution in paravermal lobules IV-V and simple lobule. Square indicates the approximate region shown in A–M. Shaded regions represent Z+ stripes in M and N. O, Analysis of PC birthdate range in stripes in stripes 1+, 2+, 3+, and 4+ of lobule V with TM9.5–TM15.5 in panels A–L.

In the PC birthdate range analysis, the birthdate distribution of PCs in stripe 4+, which started at E10.0, ranged more broadly than that of PCs in stripe 3+, which started at E12.0 (purple vs. yellow-green in Fig. 6O).

The birthdate-dependent distribution pattern of PCs in this area was summarized as follows. (1) Zebrin stripes 3- and 4- were subdivided into medial and lateral substripes that had different distributions of birthdate-specific PCs. (2) The boundary between 3- and 3b-, and the boundary between stripes 4+ and 4- were recognized as the landmark boundary (filled and open arrowheads in Fig. 6B–I). A complete switching of the complementary labeling pattern occurs at this boundary within a short period between TM12.0 and TM12.5*. In other words, stripes lateral 3- and 4+ (stripes insides the landmark boundaries) were labeled with TM 10.0–TM12.0, while the medial part of stripes 3- and other paravermal stripes medial to 3- and the medial part of stripes 4- (stripes outsides the landmark boundaries) were labeled with TM 12.5*–E13.0. (3) Lateral stripes (lateral part of 4-, 5+, 5- and 6+) mostly appeared to contain Purkinje cells of long birthdate range TM10,0–TM13.0.

### 3.7. Birthdate-dependent distribution pattern of PCs in the paramedian lobule and copula pyramidis

In the paramedian lobule and copula pyramidis, a very small number of PCs were labeled in 3-, 5+, lateral 5- and 6- with TM9.5 (Fig. 7A). Many PCs were labeled in stripe 5+ and the medially neighboring narrow area of the negative stripe in the copula pyramidis and caudal part of the paramedian lobule (Fig, 7B, C). The narrow negative area medially neighboring to stripe 5+ is named stripe e2- in the copula pyramidis. Although this area has not been named in the paramedian lobule, we also call it “e2-” in the paramedian lobule (asterisk in Fig, 7B, C). Additionally, several PCs were labeled in the lateral part of 5-, 6+, 6- and 7+ with TM10.0–TM13.0 (Fig. 7B–F). Then, labeled PCs were distributed in all stripes except for 5+ and e2- with TM12.5* (Fig. 7G). Labeled PCs were distributed in e1+ and e1- except for e2-, and in the medial part of 5-, 6+, 6- and 7+ with TM12.5 and TM13.0 (Fig. 7H, I). Nearly no PCs were labeled with TM13.5–TM15.5 (Fig. 7J, K, L).

**Figure 7.**
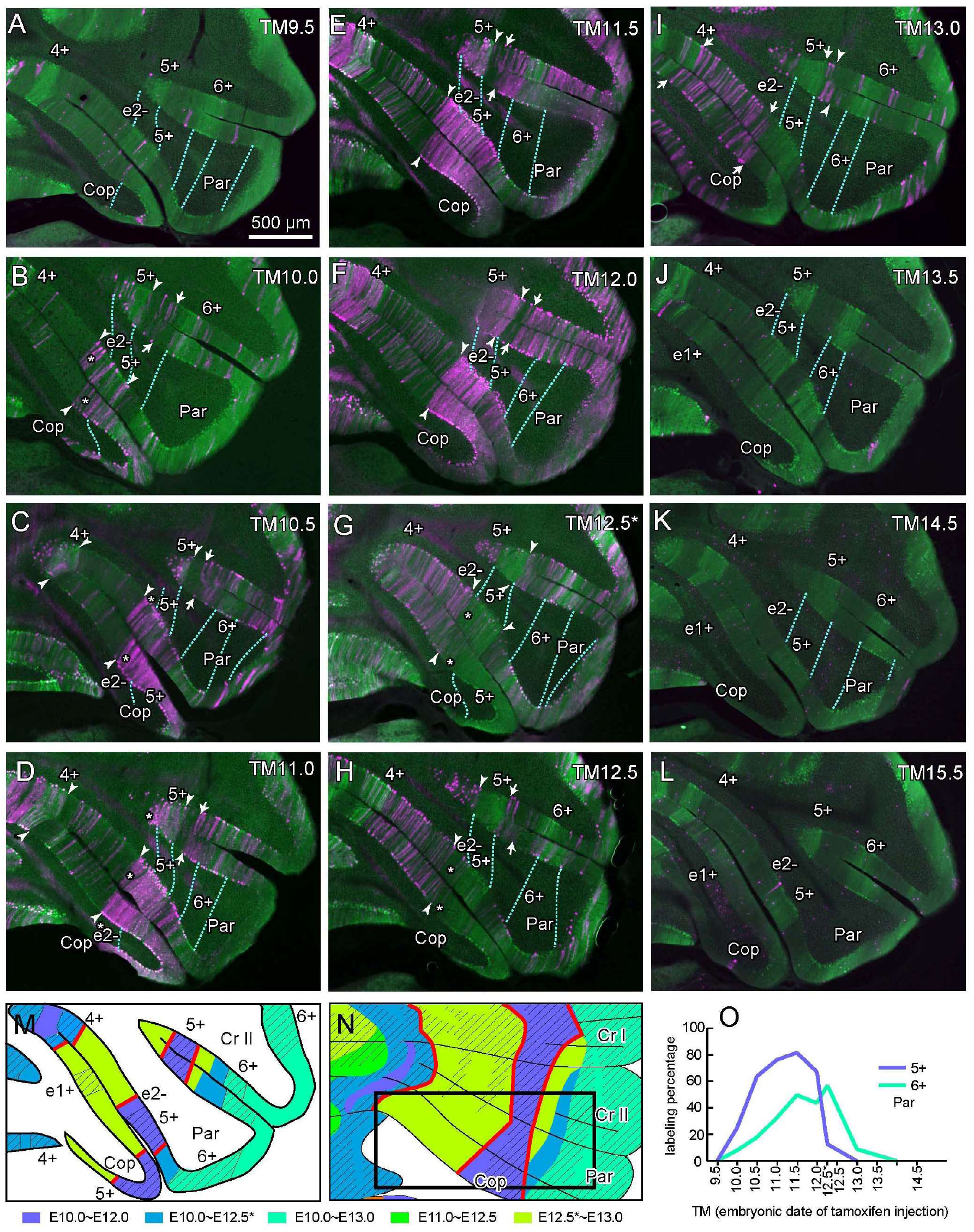
Distribution patterns of tdTomato-labeled PCs in paravermal and hemispheric crus II, paramedian lobule, and copula pyramidis in G2A::Ai9::AldocV mice at P40–42 with various tamoxifen injection timing. A-L, Images of the coronal cerebellar section at nearly the same level of G2A::Ai9::AldocV mice that were given tamoxifen at the timing indicated in the panel (E9.5–E15.5). Green and magenta channel shows the fluorescent signal of Venus and tdTomato, respectively. Arrowheads indicate the landmark boundaries of distribution of labeled PCs at the lateral edge of stripe 5+, at the medial edge of stripe 2e-, and the lateral edge of stripe 4+. Arrows indicate other medial boundaries located in variable positions. Blue dashed lines indicate the boundaries between the Z+ and Z- stripes. Scale bars, 500 µm in A–L, M. Schematic summary of PC birthdate distribution. Shaded regions represent Z+ stripes. N. Schematic summary of PC birthdate distribution in this area. Square indicates the approximate region shown in A–M. O, Analysis of PC birthdate range in stripes in stripes 5+ and 6+ of paramedian lobule with TM9.5–TM15.5 in panels A–L.

In the PC birthdate range analysis (Fig. 7O), PCs in stripe 5+ were born from TM10.0 to TM12.5*, while PCs in stripe 6+ were born in a later and longer period from TM10.5 to TM13.0.

The birthdate-dependent distribution pattern of PCs in this area was summarized as follows. (1) Stripe e2- recognized as an area of particular birthdate-dependent distribution of PCs. Stripe 5- was subdivided into medial and lateral substripes that had different distributions of birthdate-specific PCs. (2) The medial boundary of e2- was recognized as the landmark boundary (filled arrowheads in Fig. 7B–I). The boundary between stripe 5+ and 5- was also recognized as the landmark boundary (open arrowheads in Fig. 7B–I). Stripes e2- and 5+, which were circumscribed by these landmark boundaries contained TM10.0–TM12.0 PCs, whereas stripes on the other sides of these boundaries contained TM 12.5*–TM13.0. (3) The lateral part of 5- and other hemispheric stripes (6+, 6-, and 7+) mostly appeared to contain PCs born in a long period from E10.0 to E13.0.

### 3.8. Birthdate-dependent distribution pattern of PCs in the nodulus (lobule X)

No PCs were labeled with TM9.5 or TM10.0 (Fig. 8A, B). Purkinje cells were labeled in stripe 3+ and most lateral Z+ stripe, 4+, except for a narrow gap of fainter zebrin expression (3b+ in Fig. 8C-F), with TM10.5-TM12.0. Labeling in 3+ became wider with TM11.0-TM12.0. Purkinje cells were labeled in all areas except for the most lateral area with TM12.5* (Fig. 3G). PCs were labeled in medial areas with TM12.5 and TM13.0 (Fig. 8H, I). PCs were labeled only in the most medial area with TM13.5 (Fig. 8J). PCs were not labeled with TM 14.5 or TM15.5 (Fig. 8K, L).

**Figure 8.**
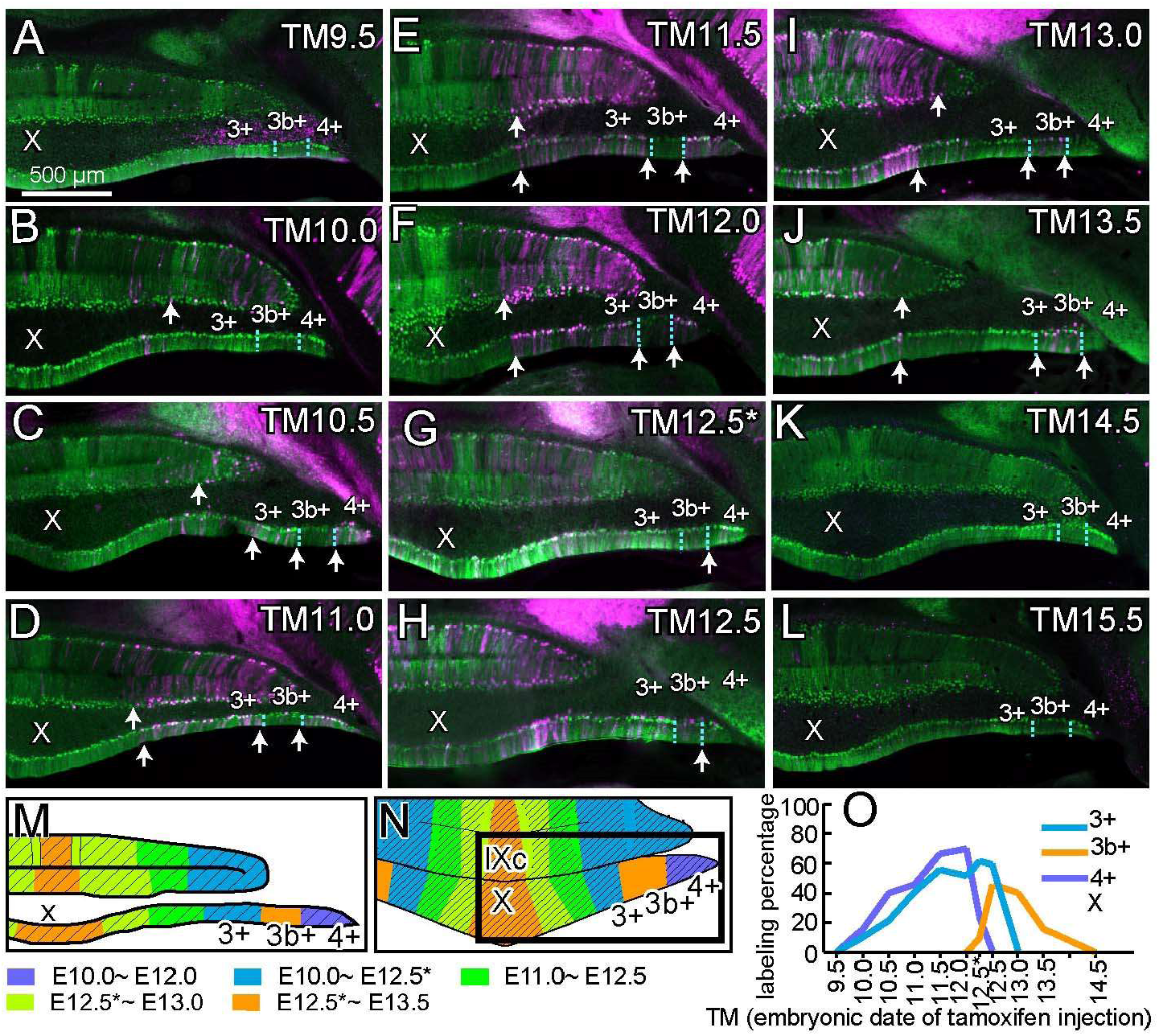
Distribution patterns of tdTomato-labeled PCs in lobule X in G2A::Ai9::AldocV mice approximately at P40–42 with various tamoxifen injection timing. A-L, Images of the coronal cerebellar section at nearly the same level of G2A::Ai9::AldocV mice that were given tamoxifen at the timing indicated in the panel (E9.5–E15.5). Green and magenta channel shows the fluorescent signal of Venus and tdTomato, respectively. Arrows indicate other medial boundaries located in variable positions. Blue dashed lines indicate the boundaries between the Z+ and Z- stripes. Scale bars, 500 µm in A–L, M. Schematic summary of PC birthdate distribution. Shaded regions represent Z+ stripes. N. Schematic summary of PC birthdate distribution in this area. Square indicates the approximate region shown in A–M. O, Analysis of PC birthdate range in stripes in stripes 3+, 3b+ and 4+.

In the PC birthdate range analysis in Z+ stripes in lobule X (Fig. 8O), PCs in stripe 4+ and 3+ were born from TM10.0 to TM12.5 or TM12.5*, while PCs in stripe 3b+ were born in a later and longer period from TM12.5* to TM13.5.

In summary, birthdate-specific PC distribution was similar to that of other vermal areas in the medial half of lobule X, but distinct in the lateral half of lobule X. In the medial half of lobule X, PC labeling shifted from the lateral to medial areas along with the shift of the labeling timing from TM10.0 to TM13.5 (Fig. 8M, N; changing positions of arrows in Fig. 8B–J). In the lateral half of lobule X, the landmark boundary of PC labeling may be present at the boundary of zebrin stripes 3b+ and 4+.

### 3.9. Birthdate-dependent distribution pattern of PCs in the paraflocculus and flocculus

Since cortical longitudinal stripes in the paraflocculus and flocculus are arranged roughly in parallel to the coronal plane (Panezai et al., 2020), horizontal sections were preferentially used to analyze birthdate-specific PC distribution in these lobules. In the paraflocculus, PCs in the caudal part (left side in panels of Fig. 9A-D) was moderately labeled with TM10.5 (Fig. 9A1–3, “p1”), densely labeled with TM11.5 (Fig. 9B1–3, “p1”), but unlabeled with TM12.5 (Fig. 9C1–3) or TM13.0 (Fig. 9D1–3). On the contrary, PCs in the rostral part were not labeled with TM10.5 (Fig. 9A1–3) or TM11.5 (Fig. 9B1–3), but densely labeled with TM12.5 (Fig. 9C1–3, “p2” and “p5”), and sparsely labeled generally with TM13.0 (Fig. 9D1–3-). In small areas in the rostral and central part PCs were densely labeled with TM13.0 (Fig. 9D2,3, “p3” and “p5”). In the flocculus, the most caudal small area had PCs labeled with TM10.5 (Fig. 9A3, “f1”). This area and another small area had PCs labeled with TM11.5 (Fig. 9B2,3, “f1” and “f3”). The rostral half of the rostral Z- part contained moderate distribution of labeled PCs with TM11.5 (Fig. 9B1,2,3, “f6”). The entire rostral Z- part contained a moderate distribution of labeled PCs with TM12.5 (Fig. 9C1,2,3, “f5” and “f6”). The caudal half of the rostral Z- part (Fig. 5 “f5”), and the neighboring small area and the separate area in the caudal Z+ part contained labeled PCs with TM13.0 (Fig. 9D2,3, “f4” and “f2”).

**Figure 9.**
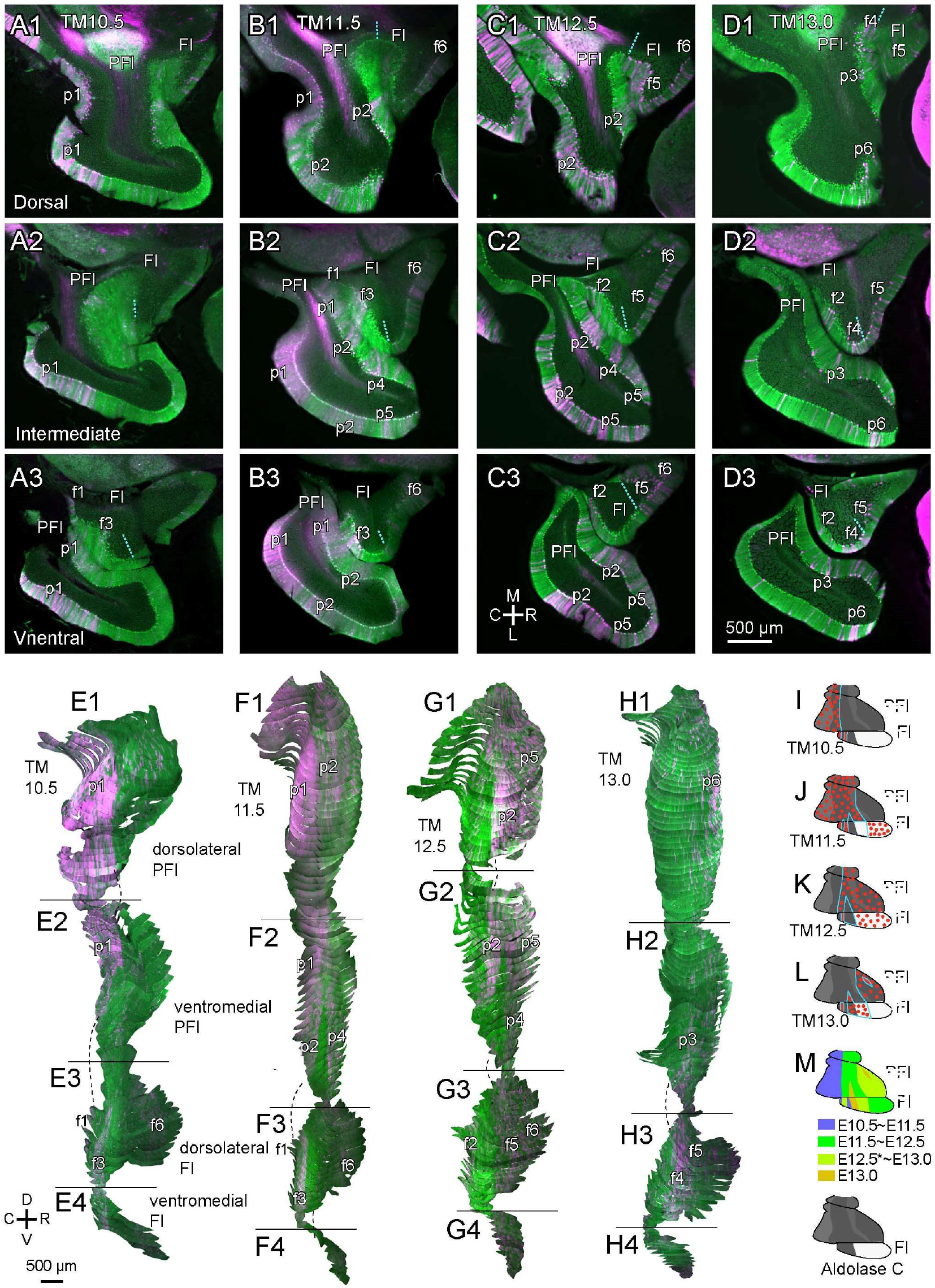
Distribution patterns of tdTomato-labeled PCs in the paraflocculus and flocculus in four G2A::Ai9::AldocV mice approximately at P40–42 with tamoxifen injection at E10.5, E11.5, E12.5, and E13.0. A–D, Images of the right paraflocculus and flocculus in horizontal cerebellar sections at three levels in the four mice. Green and magenta channel shows the fluorescent signal of Venus and tdTomato, respectively. Blue dashed lines indicate the boundaries between the Z+ and Z- stripes in the flocculus. Scale bar (in D3), 500 µm, applies to all panels. E–H. SSAA of the dorsolateral and ventromedial aspects of the paraflocculus (E1, F1, G1, H1, and E2, F2, G2, H2, respectively) and the dorsolateral and ventromedial aspects of the flocculus (E3, F3, G3, H3, and E4, F4, G4, H4, respectively) from serial horizontal sections in the four mice. Dashed lines indicate continuity of the labeling pattern across SSAA segments. Identified distribution areas of differently labeled PCs were indicated by p1- p6 in the paraflocculus and by f1-f6 in the flocculus (see section 3.9 of Results). Scale bar, 500 µm. I–L, Mapping of labeled PCs on the unfolded scheme of the paraflocculus and flocculus. M. Schematic summary of PC birthdate distribution in the paraflocculus and flocculus.

To further clarify the distribution pattern of birthdate-specific PCs, we performed serial section alignment analysis (SSAA; Sarpong et al., 2018) in serial horizontal sections of four samples with TM10.5, TM11.5, TM12.5 and TM13.0 (Fig. 9E–H). In the dorsolateral aspect of the paraflocculus, PCs were labeled moderately in the caudal part with TM10.5 (Fig. 9E1, “p1”), densely in the caudal part and moderately in the central part with TM11.5 (Fig. 9F1, “p1” and “p2”, respectively), densely in the central and rostral part with TM12.5 (Fig. 9G1, “p2” and “p5”), and sparsely in the rostral part but densely in a small localized area with TM13.0 (Fig. 9H1, “p5” and “p6”). The distribution pattern of birthdate-specific PCs in the ventral aspect of the paraflocculus was similar to that in the dorsal aspects of the paraflocculus except that the central “p2” area is divided into the caudal and rostral parts (Fig. 9F2, G2, “p2” and “p4”) with the intercalating narrow area of PCs that were labeled with TM13.0 (Fig. 9H2, “p3).

In the flocculus, labeled PCs emerged in narrow substripes inside the caudal Z+ part with TM10.5 and TM11.5 (Fig. 9E3, F3, asterisk “f1” and “f3”). Labeled PCs also emerged in the rostral subarea of the rostral Z- part with TM11.5 (Fig. 9F3, asterisk “f6”). Labeled PCs appeared in a narrow substripe in the caudal Z+ part (Fig. 9G3, “f2”) and in the entire rostral Z- part with TM12.5 (Fig. 9G3, “f5” and “f6”). They remained moderately labeled in the caudal subarea of the rostral Z- part with TM13.0 (Fig. 9H3, “f5”). Besides, Labeled PCs appeared densely in the most rostral subarea of the caudal Z+ part with TM13.0 (Fig. 9H3, “f4”). Most of these distribution areas extended into the ventromedial aspect of the paraflocculus (Fig, 9E4, F4, G4, H4). Areas in the ventromedial paraflocculus named “p1”, “p2” and “p3” appeared continuous to areas in the dorsolateral flocculus named “f1”, “f3” and “f4”.

The above-described observation in SSAA confirmed the identification of distribution areas of birthdate-specific PCs in individual sections. These areas were summarized in the unfolded scheme of the paraflocculus and flocculus (Fig. 9I–M). The distribution patterns in the flocculus and paraflocculus were different from each other and they were also different from that of the hemispheric part of other lobules. In the paraflocculus, PC distribution areas were classified into several caudal and rostral parts. (1) the most caudal part (less strong in zebrin expression) were labeled with TM10.5– TM11.5 (“p1” in Fig. 9A-H); (2) other parts were labeled with TM11.5–TM13.0 (“p2”- “p6” in Fig. 9A-H). The positional relationship between these areas and molecular compartments of the paraflocculus (Panezai et al., 2020) was not clear yet. In the flocculus, six stripes were observed with different tamoxifen timing. Noticeably, a few small areas in the paraflocculus and flocculus contained TM13.0 PCs.

### 3.10. Birthdate-dependent distribution pattern of PCs mapped on the zebrin pattern in the entire cerebellar cortex

We observed the general distribution patterns of the birthdate-specific PCs obtained with TM10.5–TM13.5 in the entire cerebellar cortex, in the way described for several particular areas in preceding sections. The results are plotted on the unfolded scheme of the zebrin striped pattern in the entire cerebellar cortex (Fig. 10A–G). TM10.5 was the earliest tamoxifen injection date when a substantial number of PCs were labeled. With TM10.5, PCs were labeled in the lateral vermis, 4+//5+ and medially neighboring Z- area, and several Z+ and Z- stripes in the hemisphere (Fig. 10A). The double slash here (4+//5+) means the combination of rostral and caudal stripes; stripe 4+ in rostral lobules (lobules II-V, simple lobule and rostral crus I) and 5+ in caudal lobules (caudal crus I, crus II, paramedian lobule and copular pyramidis) in this case (Sugihara and Shinoda, 2004). With TM11.5, the distribution of PC labeling became wider into medial stripes (Fig. 10B). With TM12.5*, the distribution of PC labeling became even wider (Fig. 10C). With TM12.5, the distribution that was nearly complementary to that with TM10.5 emerged mostly in the whole cerebellar cortex except for the hemisphere (Fig. 10D). This pattern persisted, except in a few areas, with TM13.0 (Fig. 10E). Only a few small areas were labeled with TM13.5, around the rostral c+, caudal 4b+ bands and the triangle area (above) between the paraflocculus and flocculus (Fig. 10F). Few PCs were labeled with TM9.5, TM14.5, or TM15.5 (not shown).

**Figure 10.**
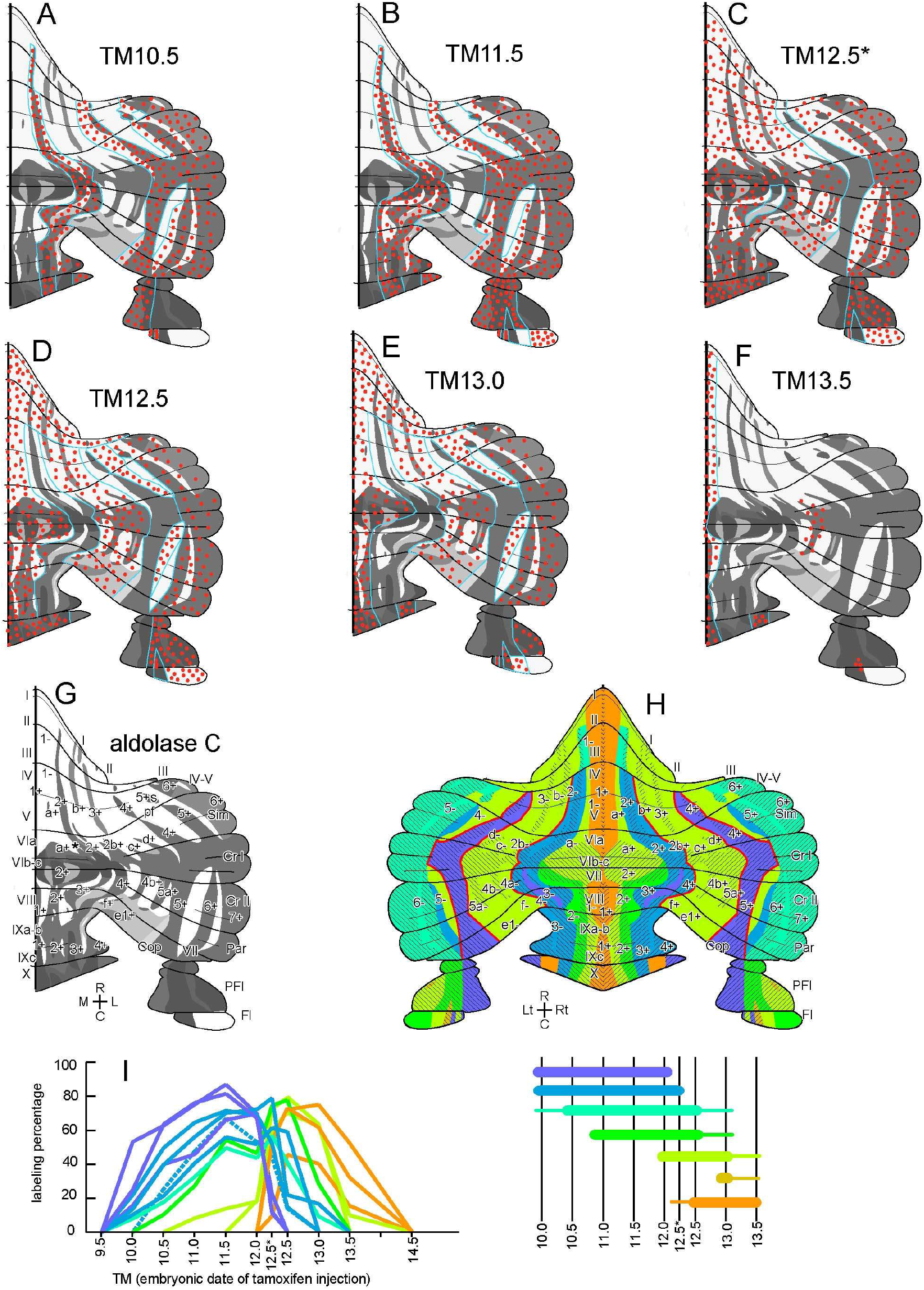
Summary of the distribution patterns of tdTomato-labeled PCs in G2A::Ai9::Aldoc mice in the whole cerebellar cortex with various tamoxifen injection timing. A–F, Distribution patterns of tdTomato-labeled PCs with tamoxifen injection at E10.5, E11.5, E12.5*, E12.5, E13.0 and E13.5. Red dots represent the distribution and expression intensity of birth-date related PCs. G, Unfolded scheme of zebrin stripes (Sarpong, 2018). H. Expression patterns of PCs born on every stage summarized in the unfolded cerebellar cortex scheme. Each color indicates different periods of tamoxifen injection effective in labeling PCs in the colored area. Thick red curves indicate landmark boundaries. I, Overlay of graphs of analysis of PC birthdate range shown in preceding figures.

As summarized in Fig. 10H, all distribution areas are arranged in longitudinal stripes. The distribution pattern was nearly consistent and continuous throughout all cerebellar lobules except for the nodulus, paraflocculus, and flocculus. The distribution patterns of birthdate-specific PCs in the flocculus and (the lateral part of) the nodulus were visibly different from those in the rest of the cerebellum. Concerning the three major conventional divisions of the cerebellum, the distribution patterns in the vermis, paravermis, and hemisphere appeared distinctly from one another. The vermis has a simple gradient organization composed of early-born PCs in the lateral part and late-born PCs in the medial part. A landmark boundary was observed at the putative lateral edge of the vermis. In the paravermis, stripe 4+//5+ and medially neighboring Z- substripe (3b-//e2-) which contained early-born PCs and the rest of stripes which contained the late-born PCs are separated by clear landmark boundaries. In the hemisphere, PCs were generally born in a long range with TM10.0–TM13.0. Consequently, no landmark boundaries were observed inside the hemisphere. This made the detailed analysis of distributions of birthdate-specific PCs more complicated in the hemisphere than in other areas. Further analysis may be required to detect detailed futures in the distribution of birthdate-specific PCs.

The superimposition of all graphs of PC birthdate range analysis in preceding sections showed various types of birthdate range among 14 Z+ stripes (Fig. 10I). Labeling percentage higher than 50% was observed in various periods between TM10.0 and TM13.5. However, the birthdate ranges of zebrin stripes seemed to be mostly classified into two major groups, a type of earlier and longer ranges (purple, blue, light blue and green in Fig. 10I) and another type of later and shorter ranges (yellow-green and orange in Fig. 10I). Almost all distribution had a tall peak of 70-85% labeling, indicating a highly efficient labeling of PCs in the birthdate tagging system with the G2A mouse line. However, a few distributions had a low peak of 45–60%, indicating lower sensitivity of labeling in some populations of PCs (see Discussion). We could not apply the PC birthdate range analysis in Z- stripes since individual PCs were not well labeled in Aldoc-Venus mice in the present study. However, types of PC birthdate range and labeling percentage in Z- stripes did not seem very different from those in Z+ stripes because neighboring Z+ and Z- stripes often show indistinguishable distributions of tdTomato-labeled PCs.

## 4. DISCUSSION

The present study mapped the distribution of birthdate-specific PCs in the entire cerebellar cortex by using the birthdate tagging system with G2A mice. PCs labeled with different timing of tamoxifen injection were distributed in different patterns of longitudinal stripes. Noticeably, PCs labeled with TM10.5–TM12.0 and those labeled with 12.5–TM13.5 were distributed in complementary patterns clearly in the paravermal area. The vermis, paravermis, hemisphere, nodulus, paraflocculus, and flocculus had different organizations of birthdate-specific PCs. The cerebellar compartmental organization revealed by this study will be discussed. We also compare this technique with other birthdate-specific labeling techniques.

### 4.1. Birthdate-specific PC labeling to classify PC populations

In the present study, birthdate-specific labeling of PCs was performed with G2A mice in which the timing of tamoxifen injection to the pregnant dam can control the birthdate-specific neuronal labeling (Hirata et al., 2019). This method is technically simpler than previously established birthdate-specific labeling methods (systemic injection of 5-bromo-2’-deoxyuridine (BrdU), systemic injection of tritium-labeled thymidine, Altman and Bayer, 1997; or ex-utero ventricular injection of replication-defective adenoviral vector, Hashimoto and Mikoshiba, 2003; Namba et al., 2011). While BrdU and 3H-thymidine label progenitor cells in S-phase at the time of injection, the adenoviral vector labels progenitor cells in M-phase during four hours after injection (Hashimoto and Mikoshiba, 2003). Neurogenin2 (Neurog2) is a bHLH transcription factor that is expressed transiently at the transition from a neural progenitor to a differentiating neuron by inducing cell cycle exit through upregulation of many downstream targets (Florio et al., 2012). Thus, expression of Neurog2 gene (*neurog2*) or CreER expression in G2A mice can mark the generation timing of a neuron (Hirata et al., 2019). Other genes that are transiently expressed at the start of differentiation, such as *ascl1* (Sudarov et al., 2011) could also be used to label PCs in a birthdate-specific way. Then, possible different characteristics among neuronal differentiation genes may need to be considered.

### 4.2. Specificity, efficiency, and timing of the birthdate-tagging recombination in G2A mice in PCs

The birthdate-tagging system with G2A mice is rather a new method to label PCs in a birthdate-dependent way. Since virtually no PCs were labeled if no tamoxifen was given and the number of labeled PCs were dependent on the dose of tamoxifen (Fig. 1), we think all PC labeling is induced by tamoxifen-induced recombination. The result that different populations of PCs were labeled by changing the timing of tamoxifen injection (Fig. 2) indicates that the labeling represents neuronal generation at the time of tamoxifen injection as designed. Then, how efficient and how precise in timing is the recombination in G2A mice? In stripes 4+ in lobule IV-V, for example, a rapid rise (0% labeling with TM9.5, 55% with TM10.0 and 60% with TM10.5, Fig. 6O), long plateau (70-85% with TM11.0-TM12.0) and rapid fall (90% with TM12.0, 5% with TM12.5*) were observed in the labeling percentage analysis. The rapid rise and fall indicate mechanisms that control rapid switching of the fate of a PC, or the compartment where it eventually belongs to, during the progress of the PC generation. In addition, the plateau with high percentage of labeling (80-90% at the plateau) observed for a long period in some areas (e.g., 70-85% labeling for a 36-hour period, E10.5–E12.0, in stripe 4+ in lobule IV-V, Fig. 4O) indicate that postmitotic CreER or *neurog2* expression persists for such a long time in some PCs. The high percentage of labeling (70–85% at the plateau) indicates the high efficiency of labeling of this system as well. However, the length and height of the plateau were variable among PCs in different stripes. For example, the plateau length was 12 hours in stripe 3+ in lobule IV-V, 36 hours in stripe 4+ in lobule IV-V (above), and 48 hours in stripe 6+ in paramedian lobule (Fig. 7O). The plateau peak was 70-85% in many stripes, whereas it was 45% in stripe 3b+ in lobule X, much lower than in other stripes (Fig 8O). Thus, the duration and efficiency of recombination seem to be variable among PC populations, which may suggest heterogeneity of PC progenitors.

### 4.3. Implication to the PC generation and the cerebellar compartmentalization

The fundamental question about the PC birthdate is how it is involved in the cerebellar organization, including the compartmentalization, topographic neuronal connection, and functional localization. For this purpose, Hashimoto and Mikoshiba (2004) applied the ex-utero injection of the replication-defective adenoviral vector into the fourth ventricle of mouse embryos to label PCs in a birthdate-specific way. Labeled PCs have been mapped on the zebrin stripes of the cerebellar cortex (Hashimoto and Mikoshiba, 2003; Namba et al., 2011). These studies have proposed the original simple idea that PCs that are generated in one of the intermittent timings at E10.5, E11.5 or E12.5, attain specific transcription mechanisms that allow them to express specific set of molecules to aggregate into a compartment and to form topographic neuronal connections (Hashimoto and Mikoshiba, 2003; Namba et al., 2011). However, we observed some complicated features in the arrangement of birthdate-specific PCs in the present analysis which were not readily explained by that idea. First, the generation of PCs is not synchronized into the same 24-hour rhythm in the entire cerebellum but it proceeds more or less in asynchronous graded timing at different regions. Second, whereas some stripes appear to contain PCs that were generated during a short period other stripes contain PCs generated during a longer period. Furthermore, PCs in neighboring Z+ and Z- stripes share the same birthdate in several places, whereas a single Z+ and Z- stripe is divided into two substripes that contained PCs of different birthdates in other places. Therefore, we think that other factors such as position and environment would also be important in determining the fate of PCs besides the birthdate. Nevertheless, the birthdate of PCs is tightly related in the compartmental organization of the cerebellum, which will be discussed for different regions of the cerebellum in succeeding sections.

### 4.4. PC birthdate organization in the vermis

In the vermis, the replication-defective adenoviral vector study observed earliest PCs (E10.5-born PCs) in stripe 2-//3-, while the present study observed the earliest PCs (TM10.5 PCs) not only in stripes 2-//3- but also in medially neighboring stripes 2+//3+ and 1-/a-//2-. This difference is presumably because of the difference in labeling timing between the two techniques. Both studies found that PCs in the median Z+ stripe (1+) and neighboring Z- stripe (1-) contains the latest PCs (E12.5-born PCs in the adenoviral study, Namba et al., 2013; TM12.5, TM13.0 and TM13.5 PCs in the present study). Both studies found that the intermediate part, zebrin stripe a+//2+, contains PCs born in the intermediate period (E11.5-born and E12.5-born PCs, adenoviral study, Namba et al., 2013; TM11.5, TM12.0 and TM12.5 PCs in the present study). The birthdate-tagging system with G2A mice seems to label PCs generated at the time range a little wider (both later and earlier) than the adenoviral vector injection in terms of the injection timing.

In summary, the results of both studies are nearly equivalent to each other in the vermis as a whole. The entire vermis had a rather simple organization of the PC birthdates. Stripe 2-//3- contains the PCs generated in the earliest period and considered to be the most lateral stripe in the vermis. PCs in the more medial stripe were generated in the later period.

In the molecular expression of zebrin (aldolase C) and protocadherin 10 (Pcdh10), and also in the olivocerebellar and corticonucelear axonal projection patterns, rather complicated compartmental organizations have been revealed in the vermis (Sugihara and Shinoda, 2004, 2007; Sugihara et al., 2009; Fujita et al., 2014; Sarpong et al., 2018). However, the PC birthdate organization of the vermis was rather simple. The mechanisms of how the simple birthdate organization in the vermis is divided into rather complicated multiple Z+ and Z- stripes (and also Pcdh10+ and Pcdh10- stripes) that contain different topographic axonal projection pattern would be an interesting question.

### 4.5. PC birthdate organization in the paravermis

In the paravermis, the adenoviral study (Namba et al., 2011) located the distribution of the earliest PCs in Z+ stripe 4+//5+. However, the present study located the distribution of the earliest PCs not only in Z+ stripe 4+//5+ but also in medially neighboring Z- stripe 3d-//e2-. The adenoviral vector method does not necessarily have high sensitivity of labeling newly-generated PCs as judged from the general sparse labeling of PCs (Namba et al., 2011). The high sensitivity (efficient recombination) of the birthdate-tagging system with G2A mice in the present study enabled precise identification of areas of birthdate-specific PC distribution. The distribution of TM10.0– TM12.0 PCs in stripes 4+//5+ and 3d-//e2- were separated by a sharp landmark boundary from the distribution of mainly TM12.5–TM13.0 PCs in the rest of zebrin stripes in the paravermis. This area that contains mainly TM12.5–TM13.0 PCs includes many Z+ and Z- stripes (lateral 1-, b+, b-, 3+, 3- in lobules II-V; 2b-, 2b+s, c+, c-, d+, d- in the simple lobule and rostral crus I; 4a-, 4b+, 4b-, 5a+, 5b- in caudal crus I, crus II and paramedian lobule; 4-, f+, f-, e1+, e1- in copula pyramidis). These zebrin stripes in the paravermis correspond to multiple modules (lateral A, B, C1, CX, C2, and C3; Sugihara and Shinoda, 2004). How this area is separated into many zebrin stripes and modules is to be clarified in terms of PC birthdate and cerebellar development in the future.

### 4.6. PC birthdate organization in the hemisphere and paraflocculus

The adenoviral study has identified early-born PCs in stripe 5-//6-. On the contrary, the present study first observed division of stripe 5-//6- into two substripes (medial and lateral substripes, 5-(med)//6-(med) and 5-(lat)//6-(lat)) that contained PCs of different birthdates. Then the present study observed PCs of a wide range of birthdates (TM10.5–TM12.5) were distributed rather randomly in multiple stripes 4- (lat)//5-(lat), 5+//6+, 5-(lat)//6-(lat), and 6+//7+, except in the medial part of stripe 5-//6- in the hemisphere. Therefore, the results of the two studies do not fit completely well.

The paraflocculus has been shown to have a longitudinal compartmental organization equivalent to that of the hemisphere (Panezai et al., 2020). However, the present results showed different distributions of birthdate-specific PCs between the hemisphere and paraflocculus. The adenoviral vector study did not fully clarify the birthday-specific organization in the paraflocculus (Namba et al., 2011). Thus, a further detailed analysis may be required in the hemispheric areas.

### 4.7. PC birthdate organization in the nodulus and flocculus

The distribution pattern of birthdate-specific PCs in the nodulus was similar to that in the vermis in the medial part, but it was different from that in the lateral part. The distribution pattern of birthdate-specific PCs in the flocculus was unique. A remaining question is how these patterns of birthdate-specific PCs match with the striped patterns of molecular expressions (zebrin=aldolase C, heat shock protein 25, protocadherin 10; Sarpong et al., 2018, Fujita et al., 2014; Schoenville, 2006) and the spatial patterns of the topographic projection of climbing fibers and Purkinje cell axons and neuronal responses (Tan et al., 1995; Ruigrok, 2003; Sugihara et al., 2004; Schoenville, 2006) in the nodulus and flocculus.

## Conflict of interest statement

Authors have no Competing Interests to declare.

## Author Contributions

All authors had full access to all the data in the study and take responsibility for the integrity of the data and accuracy of the data analysis. Study concept and design: I.S. Acquisition of data: J.Z., K.T.-A., T.H. and I.S. Analysis and interpretation of data: J.Z. and I.S. Drafting of the manuscript: J.Z. and I.S. Critical revision of the manuscript for important intellectual content: I.S.

## Acknowledgments

The authors thank Mr. Richard Nana Abankwah Owusu Mensah for proofreading the manuscript.

## Table of abbreviations including those used in Figures

1+: 1- and so on, aldolase C (zebrin) stripes 1+, 1- and so on
I–X: lobules I–X
a–c: sublobules a–c (as in IXa–b)
C: caudal
Cop: copula pyramidis
Cr I: crus I
Cr II: crus II
D: dorsal
Fl: flocculus
L: lateral
Lt: left
M: medial
Par: paramedian lobule
PFl: paraflocculus
R: rostral
Rt: right
s: satellite (as in 2b+s)
Sim: simple lobule
TM: tamoxifen injection at embryonic day, as in “TM11.5”
V: ventral

## Notes

Funding information This study was supported by Grant-in-Aid for Scientific Research (KAKENHI) from the Japan Society for the Promotion of Science (19K06919 to I.S.). K.T.-A. is a recipient of MEXT scholarship for foreign doctor course students.

### Competing Interest Statement

The authors have declared no competing interest.

